# Whole Genome and Embryo Transcriptome Analysis of Vertebrate Identifies *nxhl* Controlling Angiogenesis by Targeting VE-PTP

**DOI:** 10.1101/804609

**Authors:** Honglin Luo, Yongde Zhang, Changmian Ji, Yongzhen Zhao, Jinxia Peng, Xiuli Chen, Yin Huang, Qingyun Liu, Pingping He, Pengfei Feng, Chunling Yang, Pinyuan Wei, Haiyan Yu, Hongkun Zheng, Yong Lin, Xiaohan Chen

## Abstract

**BACKGROUND:** Angiogenesis is closely associated with angiogenesis-dependent diseases including cancers and ocular diseases. Anti-angiogenic therapeutics have been focusing on the (VEGF)/VEGFR signaling axis. However, the clinical resistance, high cost and frequent administration of anti-VEGF drugs make it urgent to discover novel angiogenic pathways.VE-PTP (*ptprb*) is a novel target with great anti-angiogenic potential. However, it is unclear whether upstream signaling pathways targeting VE-PTP exist in angiogenesis.

**METHODS:** Whole genome and embryo transcriptome sequencing were applied to discover the new gene *nxhl*. Transgenic zebrafish model, morpholino knockdown and small interfering RNA were used to explore the role of *nxhl* in angiogenesis both *in vitro* and *in vivo*. RNA pulldown, RIP and ChIRP-MS were used to identify interactions between RNA and protein.

**RESULTS:** We discovered a novel zebrafish gene *nxhl* which is a homologue of the conserved gene *nxh* that co-expressed with some key genes essential for embryo development in vertebrate. *Nxhl* deletion causes angiogenesis defects in embryo. Moreover, *nxhl* is essential to mediate effects of angiogenesis *in vivo* and *in vitro*, and *ptprb* depletion duplicates the phenotypes of *nxhl* deficiency. Importantly, *nxhl* acts upstream of *ptprb* and regulates many extreme important *ptprb*-linked angiogenic genes by targeting VE-PTP (*ptprb*) through interactions with NCL. Notably, *nxhl* deletion decreases the phosphorylation of NCL T76 and increases the acetylation of NCL K88, suggesting *nxhl* may regulate downstream VE-PTP signaling pathways by mediation of NCL posttranslational modification. This is the first description of the interaction between *nxhl* and NCL, NCL and VE-PTP (*ptprb*), uncovering a novel *nxhl*-NCL-VE-PTP signaling pathway on angiogenesis regulation.

**CONCLUSIONS:** Our study identifies *nxhl* controlling angiogenesis by targeting VE-PTP through interactions with NCL, uncovering novel upstream controllers of VE-PTP. This *nxhl*-NCL-VE-PTP pathway may be a therapeutic target in the treatment of angiogenesis-dependent diseases.

**Clinical Perspective:** *What Is New?:*

- We report a novel *nxhl-*NCL*-*VE-PTP signaling pathway that controls angiogenesis.
- We for the first time demonstrate that *nxhl* interacts with NCL which simultaneously binds to VE-PTP that plays key roles on EC adherens junction, integrity and vascular homeostasis.
- *Nxhl* also controls some other crucial VE-PTP-linked downstream angiogenic genes (such as Tie2, VEGFaa, VEGFR2, Erbb2, S1pr1 and Hey2) which explain the phenotypes induced by the *nxhl* deficiency.
- Our study indicates the key role of *nxhl* on controlling angiogenesis as an upstream regulator of VE-PTP.

*What Are the Clinical Implications?:*

- There are several ongoing researches investigating the utility of VE-PTP or NCL inhibitors on treatment of angiogenesis-dependent diseases including a range of cancers and nonneoplastic diseases, such as AMD, DME, RA and atherosclerosis.
- Targeting the *nxhl*-NCL-VE-PTP pathway may facilitate therapeutic angiogenesis in patients with cancers or ocular diseases such as DME.
- Our study highlights the great potential of *nxhl* on anti-angiogenic therapeutics by targeting VE-PTP.

---

Angiogenesis is a process of new blood-vessel spreading that is orchestrated by various angiogenic factors. It plays critical roles in reproduction, organ development and wound repair. Pathologically, it is closely related to “angiogenesis-dependent diseases” including a range of tumors and nonneoplastic diseases, such as age-related macular degeneration (AMD), ^1^ diabetic macular edema (DME), ^2^ rheumatoid arthritis (RA)^3^ and atherosclerosis.^4^ Judah Folkman suggested to consider angiogenesis as an ‘organizing principle’ in biology.^5^ This conception shifted therapeutic strategies from tumor cell-centered to anti-angiogenesis-centered.^6^ In the past decades, milestone discoveries of anti-angiogenic therapeutics have been mainly focused on the vascular endothelial growth factor (VEGF)/VEGFR signaling axis. Various inhibitors of this axis, such as Ramucirumab, have been approved for several solid cancers by FDA.^7^ Over 3000 anti-angiogenic drugs have been registered clinical trials for cancer treatment and ocular neovascular diseases,^8^ highlighting the significant value of anti-angiogenic drugs for clinical applications. However, in the clinical setting, simply blocking the existing VEGF signaling pathway or other angiogenic pathways appears to be less effective for advanced cases and often causes treatment resistance.^9^ High cost of currently used anti-VEGF drugs and their frequent dosing make new drugs targeting novel angiogenic pathways clinically necessary and highly desirable.

We specially concern the protein vascular endothelial protein tyrosine phosphatase (VE-PTP, namely *ptprb* in zebrafish) in endothelial cells (ECs), which determine the permeability and integrity of the blood vessel wall and thereby is essential for angiogenesis. VE-PTP is a member of the R3-subclass of R-PTPs and consists of 2251 amino acids with 18 domains.^10^ It is indispensable during mouse vessel development^11-13^ due to the overactivated Tie2 and increased vessel enlargement.^11, 12, 14^ Evidence shows that VE-PTP plays crucial roles in angiogenesis, EC adherens junction, integrity and vascular homeostasis.^12, 15-18^ It binds to VEGFR2, resulting in increase of VEGFR2 phosphorylation and activation.^19^ It also binds to Tie2 and negatively controls Tie2-induced vascular remodeling and angiogenesis by dephosphorylation.^14^ Suppressing VE-PTP, either by genetic deletion or specific VE-PTP inhibitor (AKB-9778 or ARP-1536) activates Tie2 and thereby regulates EC permeability, integrity and angiogenesis.^20, 21^ Its specific inhibitor AKB-9778 has been investigated in cancer^22, 23^ and retinal neovascularization,^24, 25^ such as breast cancer^26^ and DME,^28^ and has exhibited its great clinical potential. Logically, targeting the upstream genes that directly or indirectly interact with VE-PTP might be a promising strategy to overcome limitations of current anti-VEGF agents. However, few *in vivo* studies have been conducted to investigate how VE-PTP is regulated by its upstream regulators. Notably, such regulators are still unreported.

Nucleolin (NCL) is also a highly conserved gene that highly expressed in ECs. ^26, 27^ Cell surface NCL plays crucial roles in the regulation of angiogenesis and tumorigenesis via interactions with various ligands, such as VEGF,^26^ EGFR,^28^ endostatin,^29^ and HER2 (ErbB2).30 For instance, VEGF is required for NCL cell surface localization in ECs, which strengthens its contribution to the angiogenesis.^31, 32^ In addition, inhibition of cell surface NCL in ECs significantly suppresses the EC migration and prevents capillary tubule formation.^31^ Previous researches demonstrate that anti-NCL pseudopeptides N6L impairs angiogenesis both *in vitro* and *in vivo* by targeting ECs and tumor vessels.^33, 34^ Increased NCL expression is related to worse prognosis of many cancers, such as diffuse large B-cell lymphoma^35^ and pancreatic ductal cancer. ^34^ For now, a variety of aptamers or antibodies targeting NCL, such as AS1411, are under clinical investigation for anticancer treatment and demonstrating promising perspectives,^36^ highlighting its potential as a therapeutic target for anti-cancer therapy.

Similar functions of VE-PTP and NCL on angiogenesis provide us clues for further study. So far, it is unclear whether both genes closely associate and thereby mediate angiogenesis process. This should be investigated in an advanced model that facilitates *in vivo* angiogenesis assay. Zebrafish is such a valuable model system for investigation of vascular development. Experimental evidences have indicated that developmental angiogenesis in the zebrafish embryo might be an useful tool for angiogenesis research in vertebrate because of its high similarity vascular network formation and expression patterns of key genes with humans and other vertebrates. ^37, 38^ The transparency and external development of embryo and the ability to produce various transgenic germ line fish, as well as the small size and rapid development make vasculature manipulation in zebrafish feasible and more cost-effective.^39^ Conserved angiogenic signaling pathways make zebrafish as an ideal system for human angiogenesis researches and anti-angiogenic or anti-cancer drug screening.^40, 41^

Herein, we identified a novel conserved gene *nxhl*, a homologue of *nxh* which is reserved after whole genome duplication (WGD) in a teleost embryo transcriptome, by combination genome and embryo transcriptome and zebrafish model. *Nxhl* strongly controls angiogenesis both *in vitro* and *in vivo*, and acts as a critical upstream regulator of VE-PTP through interactions with NCL that binds to VE-PTP. It is a potential therapeutic target for angiogenesis-dependent diseases.

## METHODS

Materials and raw data that support the findings of this study are available upon request to the corresponding authors. A detailed description of genome sequencing associated Materials and Methods is available in the Supplemental information.

### Zebrafish Care and Maintenance

Adult wild-type AB strain zebrafish were maintained at 28.5°C on a 14 h light/10 h dark cycle. ^42^ Five to six pairs of zebrafish were set up for nature mating every time. On average, 200–300 embryos were generated. Embryos were maintained at 28.5°C in fish water (0.2% Instant Ocean Salt in deionized water). The embryos were washed and staged according to.^43^ The establishment and characterization of the *TG (zlyz:EGFP)* transgenic lines have been described elsewhere.^39, 44^ The zebrafish facility at SMOC (Shanghai Model Organisms Center, Inc.) is accredited by the Association for Assessment and Accreditation of Laboratory Animal Care (AAALAC) International.

### Zebrafish Microinjections

Gene Tools, LLC (http://www.gene-tools.com/) designed the morpholino (MO). Antisense MO (GeneTools) were microinjected into fertilized one-cell stage embryos according to standard protocols.^39^ Translation-blocking (ATG-MO) and splice-blocking morpholinos of the *nxhl* (zgc:113227, NM_001014319.2) and *ptprb* (NM_001316727.1) were designed, respectively. The standard control morpholino was used as Control MO (Gene Tools). The amount of the MOs used for injection was as follows: Control MO, ATG-MO and splice-blocking -MO, 4 ng per embryo. Effectiveness of *nxhl* and *ptprb* knockdown was confirmed by qPCR (Quantitative Real-Time PCR). For the morpholinos and primers, see Table S28.

### Zebrafish Angiogenesis Studies and Image Acquisition

To evaluate blood vessel formation in zebrafish, fertilized one-cell *fli1a-EGFP* transgenic lines embryos were injected with 4 ng *nxhl*-e1i1-MO, *nxhl*-ATG-MO, control-MO, and *ptprb*-e4i4-MO, *ptprb*-ATG-MO, control-MO, respectively. At 52 phf (*nxhl* MO) and 2 dpf (*ptprb* MO), embryos were dechorionated, anesthetized with 0.016% MS-222 (tricaine methanesulfonate, Sigma-Aldrich, St. Louis, MO). Zebrafish were then oriented on the lateral side (anterior, left; posterior, right; dorsal, top), and mounted with 3% methylcellulose in a depression slide for observation by fluorescence microscopy. The phenotypes of complete intersegmental vessels (ISVs) (i.e., the number of ISVs that connect the DA to the DLAV), parachordal vessels (PAV) and caudal vein plexus (CVP) were analyzed. Embryos and larvae were analyzed with Nikon SMZ 1500 Fluorescence microscope and subsequently photographed with digital cameras. A subset of images was adjusted for levels, brightness, contrast, hue and saturation with Adobe Photoshop 7.0 software (Adobe, San Jose, California) to visualize the expression patterns optimally. Quantitative image analyses processed using image based morphometric analysis (NIS-Elements D3.1, Japan) and ImageJ software (U.S. National Institutes of Health, Bethesda, MD, USA; http://rsbweb.nih.gov/ij/). Inverted fluorescent images were used for processing. Positive signals were defined by particle number using ImageJ. Ten animals for each treatment were quantified and the total signal per animal was averaged.

### Quantitative Real-Time PCR

Total RNA was extracted from 30 to 50 embryos per group in Trizol (Roche) according to the manufacturer’s instructions. RNA was reverse transcribed using the PrimeScript RT reagent Kit with gDNA Eraser (Takara). Quantification of gene expression was performed in triplicates using Bio-rad iQ SYBR Green Supermix (Bio-rad) with detection on the Realplex system (Eppendorf). Relative gene expression quantification was based on the comparative threshold cycle method (2–ΔΔCt) using *ef1α* as an endogenous control gene. qPCR on HUVECs were performed as similar procedures. All of the primers are listed in Table S28.

### RNA-Seq

Control MO-injected embryos and embryos injected with *nxhl* MO at 3 dpf were frozen for RNA-seq analysis. Three biological replicates of 30 embryos each were analyzed in each group. RNA was purified using RNAqueous Total RNA isolation kit (Thermo Fisher). Libraries were prepared with TruSeq RNA library Prep kit v2 (Illumina) according to the manufacturer’s protocol. Libraries were sequenced at the CCHMC Core Facility using Illumina HiSeq 2500 device (Illumina) to generate 75 bp paired-end reads. Quality of the RNA-Seq reads was checked using Fastqc [http://www.bioinformatics.babraham.ac.uk/projects/fastqc/]. All of the low-quality reads were trimmed using trimmomatic [http://www.usadellab.org/cms/?page=trimmomatic]. The trimmed RNA-Seq reads were mapped and quantified to latest Zebrafish genome assembly GRCz10 for each sample at default thresholds using RSEM [http://deweylab.github.io/RSEM/]. The mRNA levels were identified using TopHat v2.0.9 and Cufflinks and normalized by the Fragments Per Kilobase of exon model per Million mapped reads (FPKM). Differential expression was analyzed by using CSBB’s [https://github.com/skygenomics/CSBB-v1.0]. Criteria of false discovery rate (FDR) <0.01 and fold changes <0.5 or >2.0 (<–1 or >1 log2 ratio value, p value < 0.05) were used to identify differentially expressed genes. Gene Ontology (GO) annotation, domain annotation, Kyoto Encyclopedia of Genes and Genomes (KEGG) pathway annotation and enrichment were performed using ToppGene [https://topgene.cchmc.org/].

### Transwell Migration and Invasion Assays

To examine the function of human Harbi1, the homologous gene of *nxh* and *nxhl*, siRNA targeting human Harbi1 gene (NM_173811.4) was designed (see Table S28). HUVECs cells (ATCC, Manassas, Virginia, USA) were cultured in DMEM/F12 (Hyclone, USA) with 10% FBS (Gibco BRL. Co. Ltd.) and 1% penicillin-streptomycin (Sangon Biotech, China.) at 37°C in 5% CO_2_ incubator. Three experimental groups HUVECs, HUVECs + si-Harbi1 NC and HUVECs+si-Harbi1 were set and 30 pmol si-Harbi1 per well in the 24-well plates (Corning-Costa) were transferred using 9 μl Lipofectamine RNAi MAX Reagent (Invitrogen, USA). The cell migration and invasion capacity of Harbi1 on HUVECs cells were determined by transwell insert chambers (Corning, NY, USA) covered with or without 50 µl of Matrigel (1:3 dilution, BD, NJ, USA). Cells were then harvested and dissociated into a single-cell suspension. 5×10^4^ cells in serum-free medium were added to the upper chamber and 600 µl of 20% FBS-containing medium was added to the lower chamber. The chambers were then incubated for 72 h (5% CO_2_, 37 °C). Cells on the upper chamber were discarded, while cells on the lower chamber were fixed with 4% paraformaldehyde for 30 min and then stained with 0.1% crystal violet for 10 min. Cells that underwent migration or invasion were counted in triplicates in microscopic fields. The migration of nucleolin (NM_005381.3) was also examined by similar protocol above. The siRNA of NCL is listed in Tale S28.

### Tube Formation Assay

The HUVECs’ culture conditions and experimental set were identical to the transwell migration and invasion assays. Thirty pmol si-Harbi1 or si-NCL per well of the 24-well plates (Corning-Costa) were transferred by using 9 μl Lipofectamine RNAi MAX Reagent (Invitrogen, USA). Matrigel (250 μl per well, BD Biosciences, USA) was added to the plates and cultured at 37 °C for 30 min. Then, a suspension containing 5×10 ^4^ HUVECs was added to each well and cultured at 37°C in 5% CO_2_ incubator. A final concentration of 50 μM Calcein-AM (Solarbio, China) per well was added to the plates and incubated for 30 min at 37°C. Six hours later, the tube formation was observed and counted under the fluorescence microscope. The number of formed tubes represented the tube forming capability of HUVECs.

### Comprehensive Identification of RNA-binding Proteins by Mass Spectrometry (ChIRP-MS)

Zebrafish embryos (3 dpf) were collected and a total of 2 × 10^7^ cells were prepared and re-suspended in precooled PBS buffer followed by crosslinking with 3% formaldehyde for 30 min at 25°C. The reaction was stopped by incubation with 0.125 M glycine for 5 min. After centrifugation at 1,000 RCF for 3 min, the pre-binding probes (100 pmol per 2 × 10^7^ cells; probes see Table S28) were incubated with streptavidin beads for 30 min. The unbound probes were removed by washing three times. The beads with probes were incubated with the cell lysate and hybridized at 37°C overnight with shaking. All of the beads were washed 3 times with pre-warmed wash buffer for 5 min. A small aliquot (1*/*20 of the beads) of post-ChIRP beads were reserved for RNA extraction and qPCR analysis. Then 100 μL of elution buffer (12.5mM biotin, 7.5mM HEPES pH 7.5, 75mM NaCl, 1.5mM EDTA, 0.15% SDS, 0.075% sarkosyl, 0.02% Na-Deoxycholate, and 20 U benzonase) was added and the protein was eluted at 37°C for 1 h. The eluent was transferred to a fresh low-binding tube and the beads were eluted again with 100 μL of elution buffer. The two eluents were combined and the reverse-crosslinking was performed at 95°C for 30 min. The protein was precipitated with 0.1% SDC and 10% TCA by centrifugation at 4°C for 2 h. The pellets were then washed with precooled 80% acetone three times before the proteins were used for mass spectrometry (MS) analysis. Then 5 μL peptides of each sample were collected and separated by nano-UPLC easy-nLC1200 liquid phase system before they were detected using an on-line mass spectrometer (Q-Exactive) at a solution of 70,000. All of the original MS data were queried against zebrafish protein database (UNIPROT_zebrafish_2016_09). Only those proteins with an FDR < 0.01 and an adjusted *p*-value < 0.05 were considered differentially expressed. The identified proteins were then further examined using bioinformatics, including GO) and KEGG pathway annotations.

### *Nxhl* Protein Expression and Antibody Preparation

Briefly, *nxhl* gene (zgc:113227, NM_001014319.2) was synthesized and the expression plasmid pET-B2m-*nxhl*-His was constructed using the seamless cloning technology (Figure S13 and Figure S14). The plasmid was transferred into the *Escherichia coli* strain B21 (DH3) for protein expression and the resulting protein was purified by Ni-NTA chromatography column. The purified *nxhl* protein was used to immunize Japanese big ear rabbits to produce polyclonal antibody. The specificity of polyclonal antibody was detected by anti-His Western blotting and its immunity was verified by ELISA. We purified 6mg of fusion protein (62.0 kDa) with 85% purity. After immunization in rabbits,a *nxhl* polyclonal antibody with a titer of 1:256,000 was obtained. The concentration of the *nxhl* antibody purified by Protein G affinity chromatography column was 10 mg/mL and the purity was 90%. The obtained *nxhl* antibody was used to perform Western blotting assays.

### Western Blotting Assays

Zebrafish tissues from knock-down group and control were treated with 1 mL of tissue lysate (50 mmol/L Tris, 0.1% SDS,150 mmol/L NaCl, 1% NP-40, 5 mmol/L EDTA, 5 μg/mL aprotinin and 2 mmoL/L PMSF followed by lysis with protein lysate at 4°C for 30 min). All of the samples were centrifuged at 12,000 r/min at 4°C for 20 min and the supernatant was removed to detect the protein concentration using a bicinchoninic acid (BCA) kit (CWBIO. Co., Ltd., Shanghai, China). Samples were resolved by SDS-PAGE using a NuPAGE 4–12% gel (Life Technologies). Proteins were transferred onto a nitrocellulose filter (BioRad, Hercules, CA, USA) and sealed at 4°C overnight by 5% dried skimmed milk. The membranes were incubated with diluted primary rabbit polyclonal *ptprb* (VE-PTP)(PA5-68309, Invitrogen, USA) (1:1000), Hey2(PA5-72676, Invitrogen, USA) (1:2000), Dot1L(ab72454, Abcam, UK) (1:2000), S1pr1(PA5-72648, Invitrogen, USA) (1:1000), HAND2 (PA5-68502, Invitrogen, USA) (1ug/mL), Nucleolin (ab50279, Abcam, UK) (1:1000), Nucleolin (phosphor T76, ab168363, Abcam, UK) (1:1000), Nucleolin (phosphor T84, ab196338, Abcam, UK) (1:1000), Nucleolin (acetyl K88, ab196345, Abcam, UK) (1:1000), *nxhl* (Lab made, 1:1000) and *ptprb* (Lab made, 1:1000) antibodies overnight at 4°C followed by washing with PBS at room temperature. The membranes were treated with goat-anti-rabbit, rabbit-anti-goat or goat-anti-mouse IgG-HRP secondary antibody (1: 2000, CWBiotech., Ltd., Beijing, China) and incubated at 37°C for 2 h. After washing with PBS, the membrane was soaked in enhanced chemiluminescence (ECL) kit (CW Biotech., Ltd., Beijing, China) according to the manufacturer’s protocols.

### RNA Binding Protein Immunoprecipitation Assay (RIP)

To detect the interactions between *nxhl* mRNA and nucleolin protein, and VE-PTP mRNA and nucleolin protein, RIP was conducted as follows: constructed *nxhl*-pcDNA3.1 (pcDNA3.1 vector V79020, Invitrogen, USA) was transferred into 293T cells and its overexpression was verified by qPCR. Then 10^7^ 293T cells were suspended and lysed for 20 mins with 1 ml RIPA lysis buffer (Thermo Fisher Scientific, USA) containing 1 μl of protease inhibitor (Beyotime, China). Twenty μl of cell lysates were used as input, for the IgG and IP experimental group. Magnetic beads were pretreated with an anti-rabbit IgG (Beyotime, China; negative control) or anti-Nucleolin (ab50279, Abcam, UK) for 1 h at room temperature, and cell extracts were immunoprecipitated with the beads-antibody complexes at 4°C overnight.The retrieved RNA was purified by using the phenol-chloroform method and subjected to real-time qPCR and general reverse-transcription PCR for *nxhl* and VE-PTP analysis. Primers are list in Table S28.

### RNA Pull-down Assay

To detect the interactions between VE-PTP mRNA and nucleolin protein in 293T cells, and *ptprb* mRNA and nucleolin protein in zebrafish tissues, probes for VE-PTP (human) and *ptprb* (zebrafish) were designed and synthesized. Probes were list in Table S28. Probes are labeled with 3 μg biotin then heated at 95°C for 2 min followed by standing at room temperature for 30 min. Magnetic beads were washed and resuspended in 50 μl RIP buffer, then the biotinylated and denatured probes were added and incubated for 1 h at room temperature. The nucleolin protein was extracted with 1 ml RIP buffer, sonicated, centrifuged at 12,000 rpm for 15 min, and the supernatant (nucleolin protein) was retained. A magnetic separator was used to remove the liquid and the protein solution was rinsed three times using RIP buffer. The protein solution was added to the magnetic bead-probe mixture, and RNase inhibitor was added to the lysate.The mixture was incubated at room temperature for 1 h and washed five times with 1 ml RIP buffer once. Then 2×SDS loading buffer was added to the mixture, denatured at 95°C for 10 min, and used for subsequent Western blotting. The Western blotting was performed as described above. The antibody Nucleolin (ab50279, Abcam, UK) (1:1000) was used in the detection of VE-PTP (human) and *ptprb* (zebrafish) in 293T cells and zebrafish tissue.

### Statistical Analysis

All data are presented as mean ± SEM. Statistical analysis and graphical representation of the data were performed using GraphPad Prism 7.0 (GraphPad Software, San Diego, CA). Statistical evaluation was performed by using a Student’s t test, ANOVA, or χ^2^ test as appropriate. *p* value of less than 0.05 was considered statistically significant. Statistical significance is indicated by * or *p* value. * represents *p* < 0.05, and *** indicates *p* < 0.0001. The results are representative of at least three independent experiments.

## RESULTS

### WGD Drives Teleost Karyotypes Stability in Embryo

We have been thinking that whether the reserved genes after WGD in vertebrate function in regulation of angiogenesis. We tried to find such gene conserved in vertebrate. We used a teleost golden pompano which underwent WGD as an experimental model. We firstly obtained a high-quality genome by *de novo* sequencing, assembling and annotation of this teleost (Figure S1, S2, S3; TableS1-S17). The genomic landscape of genes, repetitive sequences, genome map markers, Hi-C data, and GC content of the golden pompano genome is visualized by circus ^45^ in Figure 1A. Then, we reconstructed the evolutionary history of teleost fishes with spotted gar, zebrafish and other teleosts to examine the evolutionary position of the teleost in vertebrate (Figure 1B, Figure S4). Assuming a constant rate of distribution of silent substitutions (dS)^46^ of 1.5e-8, we revealed the dates of WGD (Ts3R) and Ss4R at 350 Mya and 96 Mya, respectively (Figure 1C). Genome collinearity comparison (Figure 1D, and Table S18) implied that teleost-ancestral karyotypes are considerably conserved in post-Ts3R rediploidization with large fissions, fusions or translocations (Figure 1E, and Table S19-S20). Next, we classified the Ts3R subgenomes according to the integrity of gene as belonging to the LF, MF, and Other subgenome.^47^ The component of rediploidization-driven subgenomes is unequally distributed among subgenomes (Figure 2A), suggesting an asymmetric retention of ancestral subgenomes in teleosts.^48, 49^ For now, knowledge on the relationship between rediploidization process and embryo development stability is largely unclear. We then compared the genome-wide transcriptional levels of LF, MF, and Other karyotypes from whole-embryo development stages (OSP to YAPS) (Figure 2B). Karyotypes-retained regions (LF and MF) showed comparable expression levels during the embryo development, while karyotypes-loss regions (Other) were expressed at significantly lower levels (signed-rank sum test, *P* < 0.01) (Figure 2B, Figure S5). The Ks/Ks values of karyotypes-retained regions are significantly lower than those of karyotypes-loss regions (Figure 2C). This observation indicated that karyotypes-loss genes evolved faster than did the karyotypes-retained regions. We propose that karyotypes-retained genes are crucial for retaining embryo development stability and that karyotypes-loss genes are more prone to contribute to genetic diversity. Detail descriptions about subgenome and evolution can be found in supplementary information.

**Figure 1.**
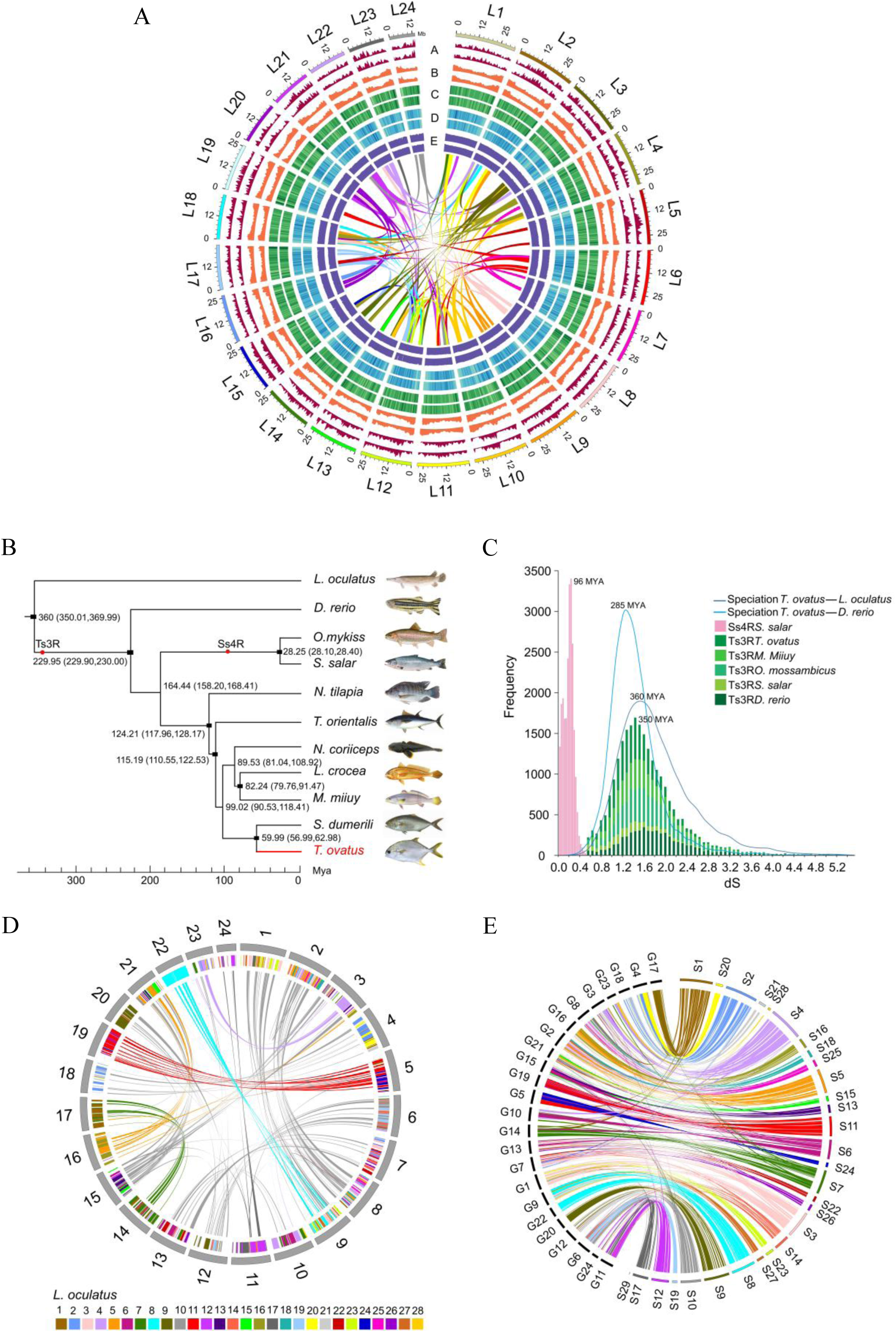
WGD of golden pompano notifies vertebrate karyotypes evolution. **A**, Overview of golden pompano (*Trachinotus ovatus)*. Numbers on the circumference are at the megabase scale. **a** Gene density of female *T. ovatus* (window size = 500 Kb). **b** TE content density of female *T. ovatus* (window size = 500 Kb). **c** Genome markers (optical) density of female *T. ovatus* (window size = 500 Kb). **d** Hi-C depth of female *T. ovatus* (window size = 500 Kb). **e** GC content of female *T. ovatus* (window size = 500 Kb). **f** Color bands in the middle of the Circos plot connect segmental duplication (minimum five gene pairs) from Teleost-specific whole genome duplication (Ts3R) events. **B**, Phylogenetic relationship of Perciformes and relevant teleost lineages. The position of golden pompano is highlighted in red. Red circles represent the Teleost specific whole genome duplication (Ts3R), Salmonid-specific whole genome duplication (Ss4R), respectively. The divergence time was estimated using the nodes with calibration times derived from the Time Tree database, which were marked by a black rectangle. **C**, Inspection of whole genome duplication events based on synonymous mutation rate (Ks) distribution. The x axis shows the synonymous distance until a Ks cut-off of 5.2. **D**, Internal genome synteny of golden pompano. Double-conserved synteny between the golden pompano and spot gar genomes. Only genes anchored to chromosomes are represented. **E**, Macro-synteny comparison between spotted gar and golden pompano shows the overall one-to-two double-conserved synteny relationship between spotted gar to a post-Ts3R teleost genome.

**Figure 2.**
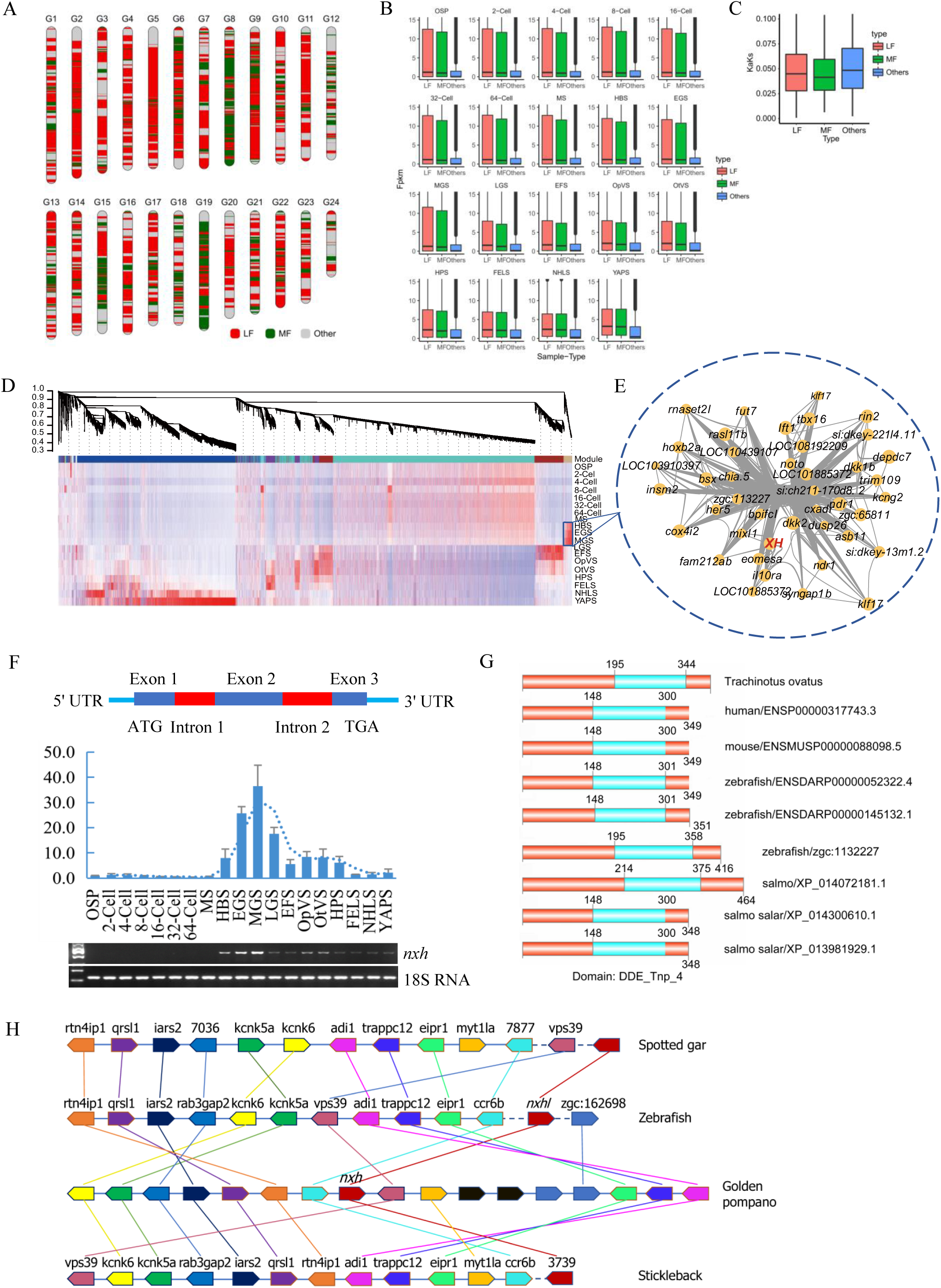
*Nxhl* is a conserved homologue of *nxh* retained after WGD. **A**, Component of less fragment (LF) and major fragment (MF) subgenomes within golden pompano genome. **B**, Boxplot of expression level of LF, MF and Other gene sets. **C**, Selection bias associated with ancestral subgenomes fragmentation. The Ka/Ks values were calculated by orthologous pairs between golden pompano and spotted gar which is outgroup species without Ts3R genome duplication events. **D**, WGCNA analysis of embryonic development stages revealed gene-network modules enriched. **E**, Hub-gene network of the purple module. Size of the dots represents hubness. Color of the dots represents the increasing expression level from low to high. Bold text highlights the genes known for *nxh* (EVM0008813) gene. **F**, Validation of expression level for *nxh* by QPCR technology. 18s RNA was considered as internal marker. Gene structure of *nxh* was showed at upper region. **G**, Micro-synteny analysis of *nxh* locus among spotted gar, zebrafish, gold pompano and stickleback. Two inversions and one insertion occurred in *nxh* locus region of golden pompano genomes. **H**, Domains of *nxh* and other homologous protein. The domains were identified in SMART database (http://smart.embl.de/).

### *Nxhl* Is A Conserved Homologue of *Nxh* Retained after WGD

Then we analyzed the gene expression pattern of golden pompano embryo (Figure S6 and Table S21-S26), and found that all 57 of the samples were separated into two components (Figure S6). The first 33 samples (from OSP to MGS) cluster into a clade and the residual 24 samples (from LGS to YAPS) cluster into another. The genes in the first clade were non-redundant reserved hub-genes and clearly “silenced” compared with those of the second clade, in which the gene levels show an explosive increase. We also noticed that before LGS, a group of genes in three stages, HBS, EGS, and MGS, are highly expressed in the first clade (Figure S6). We clustered these genes by using the WGCNA R package and found that most of them clustered into the purple_module and are co-expressed in a close network, indicating regulatory roles for these genes (Figure 2 DE). Among them, EVM0008813 (designated as New XingHuo, *nxh*; Figure 2F) is retained one copy after WGD and dominantly expresses in HBS, EGS and MGS stages. It is closely co-expressed with some key genes (Figure 2E), such as eomesa, dkk2 and mixl1, which play essential roles on embryo development.^50-52^ We purposed that *nxh* could be a crucial controller that regulates key steps of embryo development. We found that *nxh* contains 3 exons with two introns and its expression (qPCR) in EGS, MGS and LGS is highly identical to our sequencing data (FPKM, Figure 2F). We noticed that *nxh* is a WGD-specific gene and belongs to the karyotypes-retained genes (MF), implicating its important conserve function during evolution. We then searched its homologene in NCBI database by BLASTp and only one gene *zgc:113227* (designated as New XingHuo-like; *nxhl*) shares 54.7% similarity to *nxh* at the amino acid level in zebrafish. Also, the collinear analysis confirmed *nxhl* as its homologue gene in zebrafish (Figure 2G). We found that *nxh* and *nxhl* have the same functional domain DDE_Tnp_4 as the other seven genes in different species have (Figure 2H), suggesting they may have similar biological functions during embryo development. So, we asked what could the function of *nxhl* be?

### *Nxhl* Affects Angiogenic Phenotypes *In Vitro* and *In Vivo*

Firstly, we investigated whether loss of *nxhl* affects morphology development in zebrafish. We observed that both *nxhl* ^e1i1^ and *nxhl* ^ATG^ morphants resulted in nearly identical phenotypes of pericardial oedema, body axis bending, and caudal fin defects (Figure 3A, Figure S7, and Figure S8) at 3 days post fertilization (dpf), confirming that the phenotype of *nxhl* knockdown is *nxhl*-specific (Figure 3A). Regarding the vascular system, embryos injected with *nxhl* ^e1i1^ MO present thinner ISVs (yellow arrows) and ectopic sprouts (asterisk) of dorsal aorta compared with controls, and the *nxhl* knockdown prevents the parachordal vessel (PAV) formation, the precursor to the lymphatic system. Moreover, heartbeat and circulation in the caudal vein (CV) is visible in the control fish, but is abnormal in *nxhl*-MO-injected fish (Supplementary Movie1, 2). Both *nxhl* ^e1i1^ and *nxhl* ^ATG^ morphants dramatically disrupted normal splicing of *nxhl* (Figure 3A), indicating high efficiency and specificity of the morpholino knockdown of *nxhl*. Consistent with this, *nxhl* morphants resulted in a high percentage of embryos with defects (81.55%, n=103 embryos in *nxhl* ^e1i1^ MO and 100%, n=106 embryos in *nxhl* ^ATG^ MO) and low survival rate (45.78%, n=225 embryos in *nxhl* ^e1i1^ MO and 17.68%, n=198 embryos in *nxhl* ^ATG^ MO) compared with controls (n=218 embryos) at 3 dpf (Figure 3A and Figure S7B). This confirmed that knockdown of *nxhl* certainly causes morphological defects in the heart and caudal fin in zebrafish.

**Figure 3.**
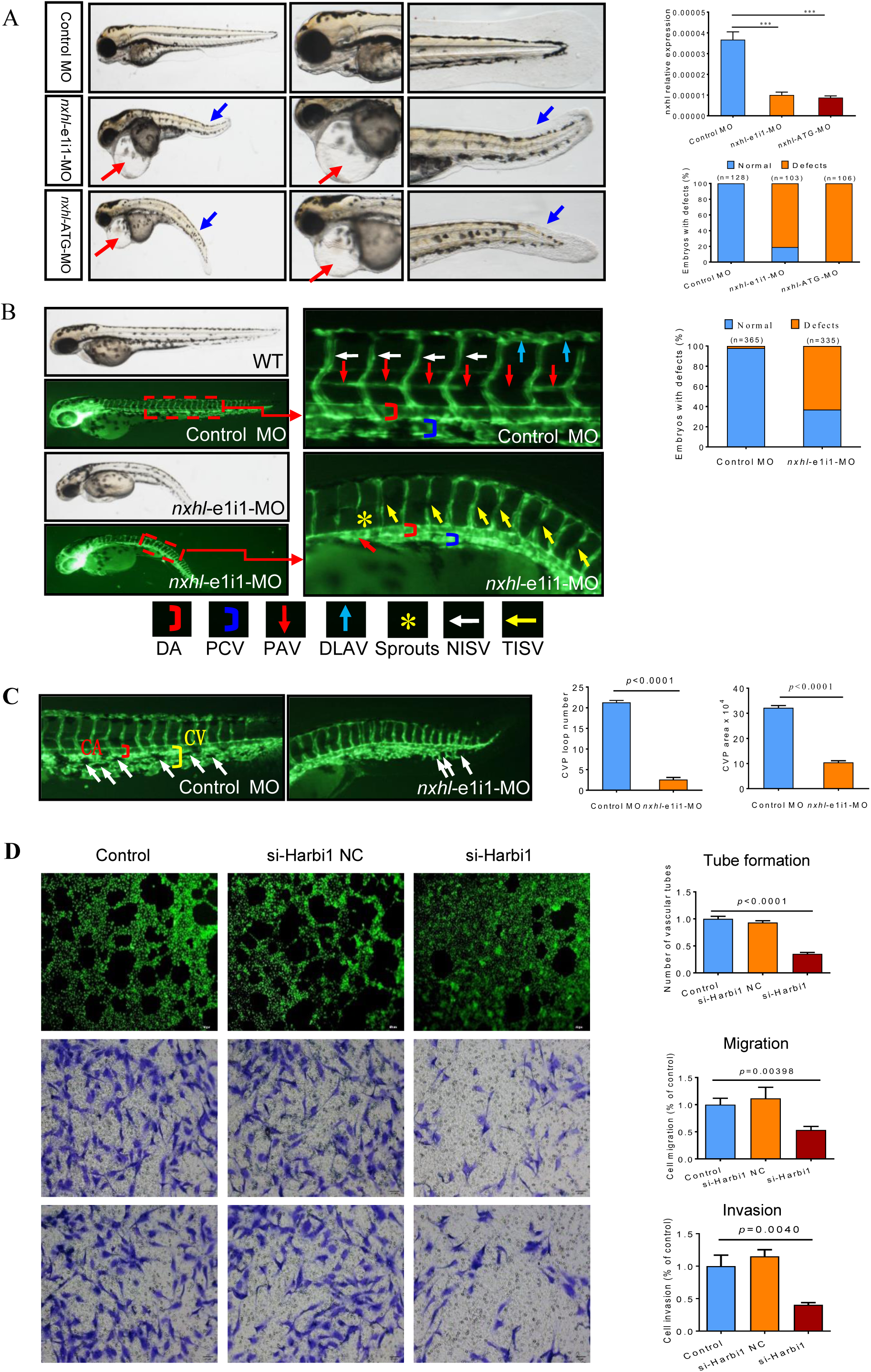
*Nxhl* affects angiogenic phenotypes *in vivo* and *in vitro*. **A**, Gross morphology at 3 dpf in wild-type AB strain. Knock down *nxhl* present pericardial oedema (red arrow) and caudal fin defects (blue arrow). The bar graph shows the validation of MO against *nxhl*, and the percentage of embryos with development defects after knockdown of *nxhl* with e1i1-MO and ATG-MO. **B**, knockdown of *nxhl* causes angiogenic defects in *Tg(fli1a:EGFP)*^*y1*^ zebrafish. Images represent bright field and fluorescent filed of *Tg(fli1a:EGFP)*^*y1*^ embryos at 52 hpf, with the angiogenic structures visualized by GFP fluorescence and labelled ISV and DLAV. The bar graph shows the percentage of embryos with angiogenic defects after knockdown of *nxhl* with *nxhl-*e1i1-MO. **C**, *nxhl* knockdown impairs formation of the CVP in zebrafish. Quantification of loop formation and area at CVP at 52 hpf. CA, caudal artery; CV, caudal vein. NISV, normal intersegmental vessel; TISV, thinner intersegmental vessel. **D**, Silence of Harbi1 inhibits angiogenic development *in vitro*. The tube formation, cell migration and invasion potential of HUVECs treated with si-Harbi1 was determined by using transwell chambers as described in the “Materials and methods” section. Scale bars, 50 μm. Representative images of cells stained in si-Harbi1 treated HUVEC cells. The data represent as mean±SEM from three independent experiments. \**p*<0.05 *p*<0.05, \*\**p*< 0.001 represents statistically significant.

We then used the *Tg(fli1a:EGFP)*^*y1*^ zebrafish as a model to investigate the connections between the vascular system and these phenotypes. Embryos were injected with 4 ng control MO or *nxhl* ^e1i1^ MO. We found that loss of *nxhl* caused intersegmental vessel (ISV) growth defect and disruption of the honeycomb structure in the CVP at 52 hpf (Figure 3B). Also, *nxhl* ^e1i1^ morphant resulted in a thinner ISV growth and ectopic sprouts of dorsal aorta at the rear-somite with only 10% of complete ISVs (n=365 embryos) compared with 98% of complete ISVs in controls (n=335 embryos). We observed that the *nxhl* knockdown prevented the PAV formation (Figure 3B) and caused specific defects in CVP formation (Figure 3C). Quantification of loop formation and the area at CVP showed a 8.2-fold and 3-fold decrease in *nxhl* ^e1i1^ morphants (n=10 embryos) at 52 hpf, respectively (Figure 3C). Our data indicate that *nxhl* plays a critical role in controlling PAV, ISV and CVP formation and vascular integrity during angiogenesis, which is an explanation strongly consistent with the heart and caudal fin phenotypes observed. What is the mechanism behind?

Endothelial cells (ECs) line the inner lumen of vessels and are the building elements of blood vessels, we then speculated that *nxhl* may affect angiogenesis via ECs. This is supported by the significant enrichment of the genes involved in blood vessel morphogenesis when *nxhl* was knocked down (Table S29). Since human Harbi1 gene shares DDE_Tnp_4 domain with *nxhl* (Figure 2E), it is supposed that both genes play similar roles. We used human umbilical vein endothelial cells (HUVECs) as a vascular epithelioid cells model *in vitro*. Next, we designed siRNAs targeting human Harbi1, transferred siRNA into HUVECs and investigated their cell migration, invasion, and tube formation. Silence of Harbi1 significantly inhibited the tube formation and cell migration compared with controls (Figure 3D). Furthermore, the invasion abilities in Harbi1 defect cells are also significantly inhibited compared with controls (Figure 3D). Moreover, silence of Harbi1 significantly inhibited the angiogenesis of non-small cell lung cancer cell (A548) and human colon cancer cell (HCT116) *in vitro* (Figure S9). This highlights the pro-angiogenesis function of Harbi1 and indicates that *nxhl* like their human homolog Harbi1, play role in angiogenesis and anti-cancer process via ECs.

### *Nxhl* Regulates *Ptprb* Expression and Angiogenic Networks

To investigate how *nxhl* mediates angiogenesis, we firstly examined transcriptome sequencing (RNA-seq) data from zebrafish after injection of 4 ng *nxhl* ^e1i1^ MO at 3 dpf. We found that loss of *nxhl* greatly changes the transcriptome with 1955 down-regulated and 698 up-regulated (Figure S10; Table S21). We noticed that in the KEGG pathways associated with angiogenesis development are significantly enriched in the *nxhl*-silenced group (Figure S11; Table S22-26). We speculated that the transcription of genes linked to angiogenesis development may also be significantly changed in the *nxhl*-silenced zebrafish. We then screened and examined the expression of 18 genes that previously documented to be closely related to heart defects and/ or angiogenesis.^53-57^ Consistent with the RNA-seq data, we found that 13 of these genes (*ptprb, tie2, nr2f1a, s1pr1, hey2, dot1L, hand2, erbb2, klf2a, mef2cb, mef2aa, ephB2a* and *cx40*.8) were significantly decreased while two genes (*vegfaa and vegfr2*) increased sharply. *S1pr2, egfl7, and nrg2a* were kept unchanged (Figure 4A-D). Notably, the arterial marker *ephB2a* and venous marker *erbb2* were decreased in *nxhl* morphants compared to the wild-type (Figure 4D). Normally, the increase of v*egfaa and vegfr2* is linked to the enhancement of vascular system.^58, 59^ However, in our study, both genes increased while others decreased when *nxhl* was silenced. We speculated this is a consequence of a negative feedback regulation to avoid an excessive decrease in the vascular system. We found that *nxhl* ^e1i1^ morphants result in decrease of the *nxhl* at protein level. *Ptprb*, the most decreased gene at mRNA level, is also greatly reduced at the protein level. The s1pr1, hand2, dot1L, and hey2 proteins were also downregulated compared with controls (Figure 4E). As previously reported, *ptprb, tie2, nr2f1a, s1pr1, vegfaa* and *vegfr2* normally contribute to vascular development and deletion of each of them leads to defects on the vascular system during embryo development,^14, 58-61^ while loss of *dot1L, hand2, erbb2, mef2cb, mef2aa, ephB2a* or *cx40*.*8* always results in angiogenesis system or heart development defects.^55, 62-68^ *Hey2 and klf2a* have been implicated in the regulation of both angiogenesis and heart development.^69, 70^ Based on these reports, we built a schematic diagram of the network as shown in Figure 4F. This network demonstrates that silence of *nxhl* does downregulate the key genes that are essential for heart and /or vascular development. To this end, our results showed that loss of *nxhl* greatly affects the expression of these key genes in the network, suggesting that the heart and vascular phenotypes caused by *nxhl* deletion are greatly due to the regulation of these genes, and the expression profiles of these genes explain the *nxhl* deficient-induced phenotypes. Thus, we then asked how *nxhl* controls the angiogenesis and angiogenic networks.

**Figure 4.**
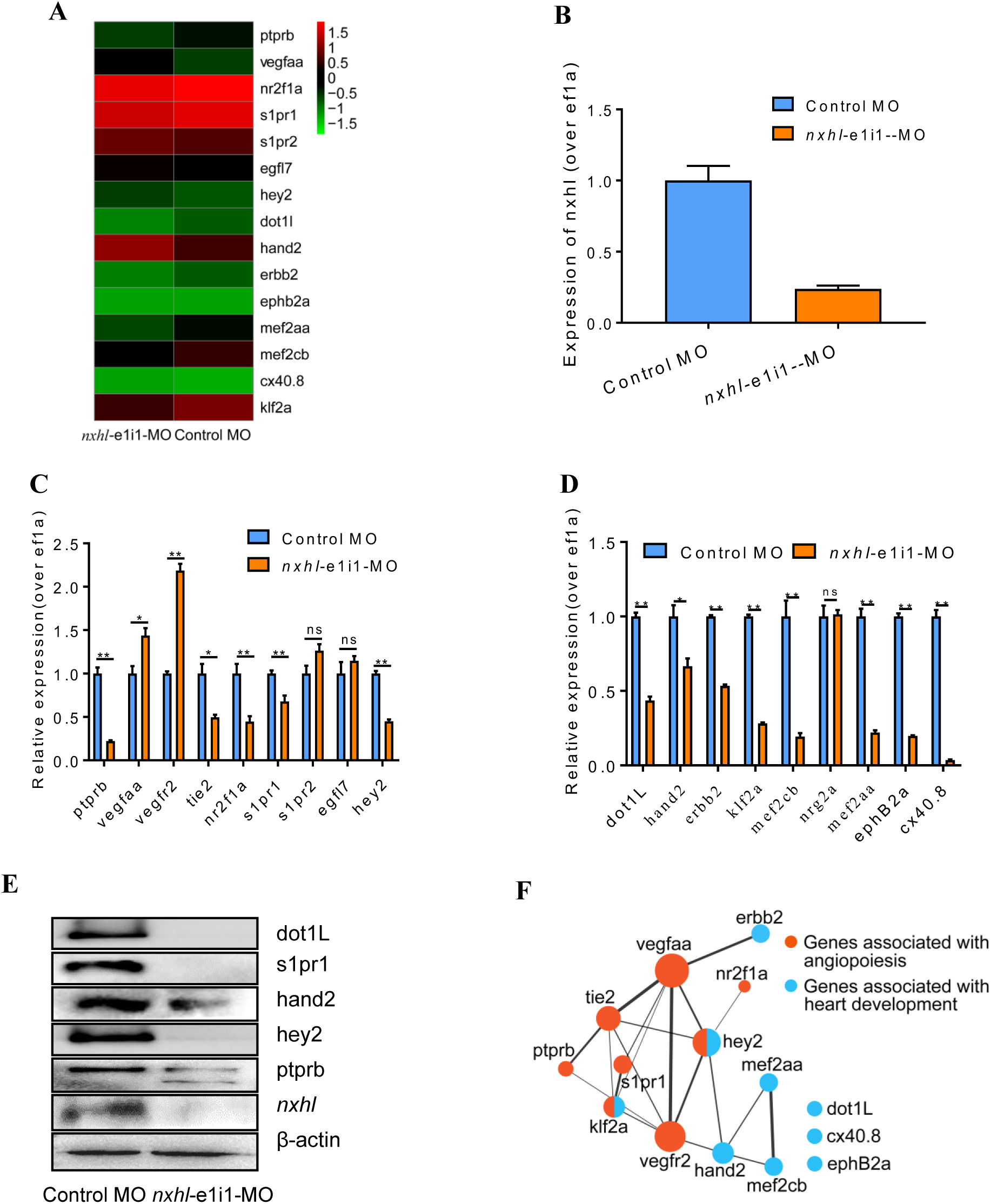
*Nxhl* modulates *ptprb* expression and angiogenic networks. **A**, Heatmap of the 15 selected genes from zebrafishes after injection of 4ng *nxhl* ^e1i1^ MO at 3 dpf examined by RNA-seq. **B**, Expression of *nxhl* post injection of *nxhl* ^e1i1^ MO 3 dpf. **C**, Expression of genes associated with angiopoiesis post injection of *nxhl* ^e1i1^ MO 3 dpf using QPCR. **D**, Expression of genes associated with heart development post injection of *nxhl* ^e1i1^ MO 3 dpf using QPCR. **E**, Networks of the genes previously reported to be associated with angiopoiesis and heart development. Cytoscope V3.6.1 was used to build this network. **F**, Protein levels of the selected genes associated with angiopoiesis and heart development post injection of *nxhl* ^e1i1^ MO 3 dpf by using Western blotting. β-actin antibody was used as internal control. The data above represent as mean±S EM from three independent experiments. \**p*<0.05 *p*<0.05, \*\**p*< 0.001 represents statistically significant.

### Loss of *Ptprb* Duplicates the Phenotypes of *Nxhl* Deficiency

As described above, we noticed that *ptprb* is the most downregulated gene after silence of *nxhl* and is the one that closely linked to both vascular integrity and angiogenesis as well.^11, 13, 19, 71-73^ To test whether there is a positive connection between *nxhl* and *ptprb*, we silenced *ptprb* by injection of 4 ng *ptprb* ^*e4i4*^ and *ptprb* ^*ATG*^ morphants designed (Figure S12, Table S30). Both *ptprb* morphants resulted in slight pericardial edema, shortened body axis and severe body axis bending in zebrafish (Figure 5A;Figure S12). Moreover, heartbeat and circulation in the caudal vein (CV) is visible in the control fish (Supplementary Movie 3,4), but is abnormal in *ptprb*-MO-injected fish (Supplementary Movie 5-8). *Ptprb* morphants also resulted in a high percentage of embryos with defects (75.48%, n=208 embryos in *ptprb* MO and 0.94%, n=212 embryos in control) and lower survival rate compared with controls at 50 hpf (Figure 5A). Both *ptprb* morphants dramatically disrupted normal splicing of *ptprb* (Figure 5B-D), decreased the survival rate, but unchanged the *nxhl* expression, indicating high efficiency and specificity of the morpholino knockdown of *ptprb*. In the vascular system, loss of *ptprb* leads to an indefinite absence or deformity of DLAVs (blue arrowhead) and ISVs in the tail end (white and yellow arrowhead), and a decrease of PAV (red arrowhead) formation (Figure 5E). Knockdown of *ptprb* caused significantly decrease of the mean diameter of ISVs compared with controls (Figure 5F). Also, *ptprb* morphants caused CVP sinus cavities defects (Figure 5G), and resulted in a 5.6-fold and 2.2-fold decrease of CVP loop formation and CVP area (n=83 embryos) at 50 hpf, respectively (Figure 5G). These data are to some extent consistent with previous reports^57^ and strongly suggest that loss of *ptprb* phenocopies *nxhl* deficiency. Moreover, We found that most of the 15 genes in *ptprb*-knockdown experiment present an expression profile similar to that in *nxhl*-knockdown experiment, except vegfaa and vegfr2 (Figure 5H). To this end, we logically concluded that knockdown of *ptprb* mimics phenotypes of *nxhl* deficiency, and both should act in the same signaling pathway. However, which one is downstream of the other is unclear. Therefore, we examined the *nxhl* expression after silence of *ptprb*, and we found that it was kept unchanged (Figure 5D), but we observed a significant decrease of *ptprb* expression after silence of *nxhl*. This confirms that *ptprb* acts at the downstream of *nxhl*.

**Figure 5.**
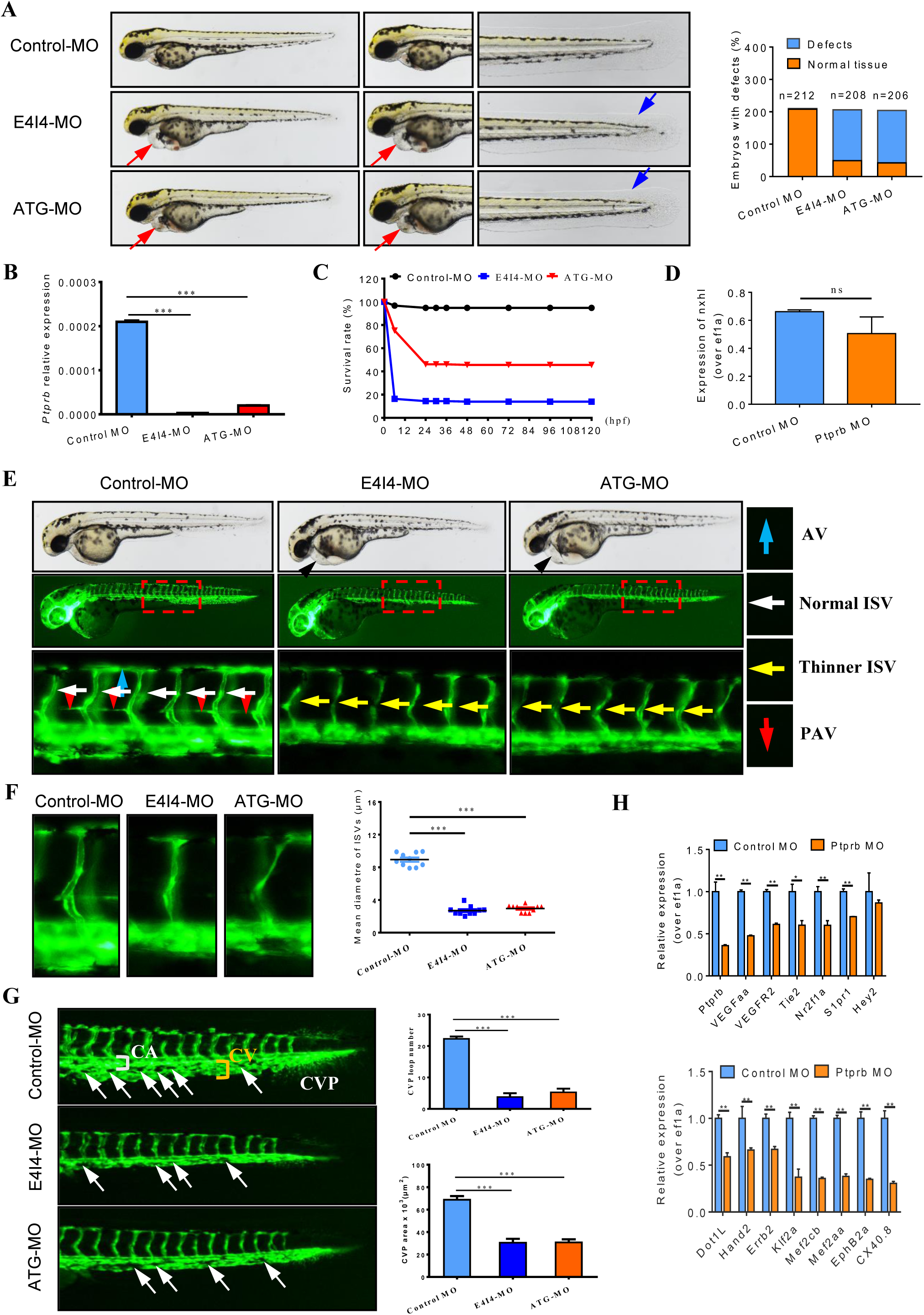
Loss of *ptprb* phenocopies *nxhl* deficiency. **A**, Gross morphology at 3 dpf. Knock down *ptprb* present pericardial oedema (red arrow) and caudal fin defects (blue arrow). The bar graph shows the percentage of embryos with development defects after knockdown of *ptprb*. **B**, Endogenous *ptprb* in control and *ptprb* morphants were assessed by qPCR. **C**, A time-course plot of percent survival in control and *ptprb* morphants for 3 days. dpf, days post fertilization. **D**, Expression of *nxhl* post injection of *ptprb* MO 3 dpf. **E**, Morpholino knockdown of *ptprb* causes angiogenic defects. Representative bright field and fluorescent images of *Tg(fli1a:EGFP)*^*y1*^ embryos at 50 hpf with the vascular structures visualized by eGFP fluorescence and labelled ISV and DLAV. The boxed regions are shown at higher magnification in the bottom panels. **F**, Quantification of the mean diametre of ISVs shows significantly decrease in *ptprb*-MO injected embryos. Columns, mean; SEM (n =10; ANOVA;) DLAV, dorsal longitudinal anastomotic vessels; ISV, intersegmental vessel. **G**, *ptprb* knockdown impairs formation of the CVP in zebrafish. Bars show the quantification of loop formation and area at CVP. CA, caudal artery; CV, caudal vein. **H**, Expression of genes associated with angiopoiesis (above) and heart development (down) post injection of *ptprb* MO 50 hpf using QPCR. The data represent as mean±S EM from three independent experiments. \**p*<0.05 *p*<0.05, \*\**p*< 0.001 represents statistically significant.

### *Nxhl* Regulates VE-PTP (*ptprb*) through Interactions with NCL

As mentioned above, among the 18 genes associated with heart and vascular development, 15 genes were significantly changed by both *nxhl* and *ptprb* morphants. We suppose these genes may be part of a regulatory network of their own. We then built a schematic diagram of the network according to previous reports (Figure 4F). This network presents connections between most of these genes, suggesting a cooperative regulation mechanism on the heart and vascular development. As we already knew that *ptprb* acts downstream of *nxhl*, we next asked if *nxhl* directly interacts with *ptprb* to mediate these genes. Thus, we designed *nxhl* probes and conducted a ChIRP-MS experiment in zebrafish to find out those proteins binding to *nxhl*. Eleven proteins with change folds above 2 were discovered (Figure 6A). This indicates that the *nxhl* RNA may interact with these proteins. Unexpectedly, *ptprb* was not found in these proteins (Figure 6A). This suggests that proteins other than *ptprb* may interact with *nxhl*. We next focused on the proteins that are associated with the vascular system, and nucleolin (NCL) (Figure 6A) aroused our interest because of its molecular conservation and important functions on angiogenesis.^31, 74^ Loss of NCL in zebrafish causes oedema and body axis bending,^75^ as well as suppression of adhesion, proliferation and migration of HUVECs.^76^ These phenotypes are identical to the phenotypes caused by *nxhl* depletion, suggesting NCL may associate with *nxhl*. To figure out whether *nxhl* interacts with NCL, we performed RNA Immunoprecipitation (RIP) using the NCL protein as bait protein in 293T cells (Figure S13, S14, S15) and then detected the *nxhl* RNA using qPCR. We found that *nxhl* RNA is significantly higher than that in IgG control in the RNAs pulled-down by the NCL protein (Figure 6B). The RNA pulled down was amplified and the sequencing results confirmed that it is *nxhl* mRNA. This indicates that the NCL protein reversely interacts with *nxhl* RNA. Therefore, these experiments prove that *nxhl* RNA and NCL protein interact physically.

**Figure 6.**
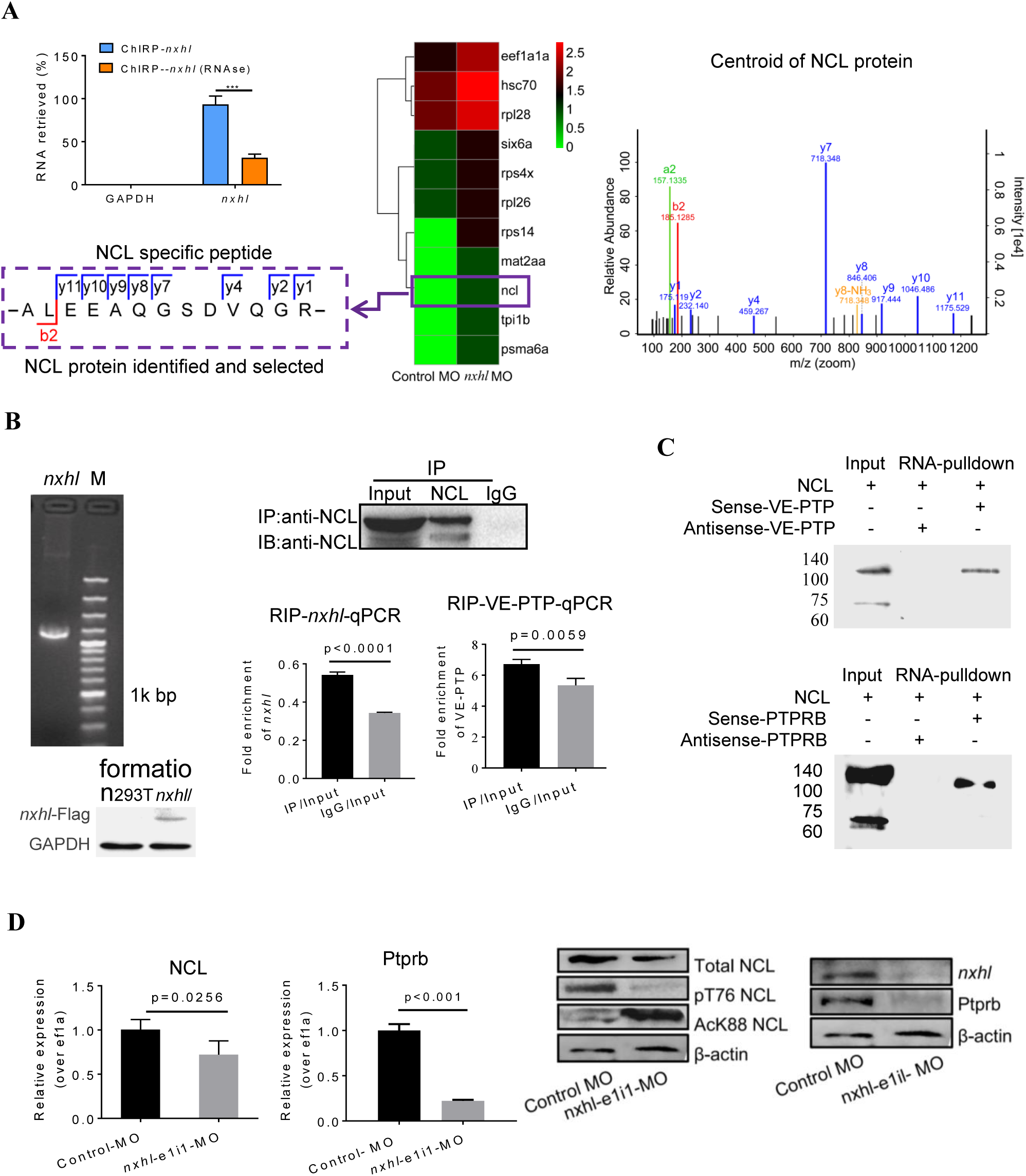
*Nxhl* regulates VE-PTP (*ptprb*) through interactions with NCL. **A**, ChIRP-MS identification of *nxhl* RNA binding proteins. qPCR identification of *nxhl* RNA in the eluted RNAs. Graph shows more than 90% *nxhl* RNA was retrieved, and no GAPDH was detected. Heat map shows major proteins are enriched and significantly (change fold >2 and p<0.05) retrieved by *nxhl* and control probes, analyzed by LC/MS-MS. NCL protein (purple boxed) was selected as candidate for follow-up study. The Centroid of NCL protein shows that NCL protein is pull down and identified by LC/MS-MS. The specific peptide identifies NCL protein. **B**, RIP-qPCR assay to detect the interaction between *nxhl*, VE-PTP mRNA and NCL protein. The mRNA expression of *nxhl* was determined by qPCR and Western blotting against Flag antibody was performed to identify the successful expression of pcDNA3.1-Flag-*nxhl* plasmid in 293T cells. Bars show the interaction between *nxhl* mRNA and NCL protein. The interaction between VE-PTP mRNA and NCL protein is shown too, and qPCR shows the detection for VE-PTP mRNA expression in the NCL-pulled down RNA. **C**, Pull down assay to detect the interaction between *nxhl*, VE-PTP mRNA and NCL protein. Gels show the interaction between VE-PTP mRNA and NCL protein. Western blotting was performed to detect NCL protein in the VE-PTP-biotin probe -pulled down proteins in 293T cells. The interaction between *ptprb* mRNA and NCL protein is shown too. **D**, Loss of *nxhl* affects the expression of NCL at both mRNA and protein levels. The mRNA expression of NCL and *ptpr* were determined by qPCR. The total NCL protein, phosphorylated NCL, acetylated NCL, total *nxhl* and *ptprb* protein were detected by Western blotting using specific NCL antibodies. The mRNA expression of *ptprb* was determined by qPCR. The data represent as mean±SEM from three independent experiments. \**p*<0.05, *p*<0.05, \*\**p*< 0.001 represents statistically significant.

However, still no evidence was found on the interaction between *nxhl* and *ptprb*. Could it be that NCL interacts with *ptprb*, thus bridging *nxhl* and *ptprb?* Such scenario was never proposed or documented before. However, a report showed that VEGF interacts with NCL.^77^ As *nxhl* acts similarly to VEGF on angiogenesis development, we then supposed that NCL might also interact with *ptprb* or its human homologue VE-PTP, the key molecule in angiogenesis. To test this hypothesis, we detected VE-PTP mRNA using the same RNAs pulled-down by NCL protein, and we found that VE-PTP mRNA is significantly higher than that in IgG control. The RNA pulled down was amplified and the sequencing results proved that it is VE-PTP mRNA. This confirms that the NCL protein can also interact with VE-PTP mRNA physically (Figure 6B). We next verified this interaction in 293T cells using the VE-PTP RNA pulldown experiment in the reverse way, and the result of western blotting against NCL protein supports the existence of interaction between VE-PTP and NCL (Figure 6C). However, whether this interaction occurs between NCL and *ptprb* in zebrafish is unclear. We next designed a zebrafish *ptprb* gene-specific probe to pull down the proteins that interact with *ptprb* in the juvenile zebrafish. We found that the NCL protein strongly binds with *ptprb* (Figure 6C). These results indicate that the NCL protein not only interacts with VEP-PTP in 293T cells but also with *ptprb* in zebrafish.

So far, we proved that *nxhl* and NCL, NCL and VEP-PTP (*ptprb*) interact physically. However, how *nxhl* regulates NCL and *ptprb* is unclear. To address this issue, we micro-injected 4 ng *nxhl*-e1i1-MO in one cell stage embryo, and found that resemble phenotypes were induced as that shown in Figure 3AB and Figure S16. Meanwhile, we found that loss of *nxhl* not only causes a significant decrease of NCL mRNA and total protein level but also leads to decrease of phosphorylated T76 and increase of the acetylated K88 of the NCL protein (Figure 6D). This suggests that knockdown of *nxhl* significantly affects the expression of NCL, which plays vital functions in angiogenesis,^31^ although the impact of phosphorylation and acetylation of NCL protein on the heart and vascular development have not been deeply understood yet.^74^ Then we investigated the expression of the downstream gene *ptprb*, and found that loss of *nxhl* also decreases *ptprb* at both mRNA and protein levels (Figure 6D). These results suggest that silence of *nxhl* leads to angiogenesis defects due to the downregulation of both NCL and *ptprb* via the interactions of *nxhl-*NCL and NCL*-ptprb*, which consequently mediates the angiogenesis-linked landmark gene network.

### NCL Regulates Angiogenesis and VE-PTP *in vitro*

Although the physical interactions between *nxhl* and NCL and NCL and *ptprb* (VE-PTP) and regulatory role of *nxhl* on NCL and *ptprb* (VE-PTP) are confirmed in our study, whether NCL regulates *ptprb* (VE-PTP) is still unknown. We next examined the functions of NCL on angiogenesis and expression of VE-PTP by silence of NCL in HUVECs. As shown in Figure 7, silence of NCL not only significantly inhibited the tube formation but also the cell migration of HUVECs comparing with the controls. Notably, silence of NCL greatly decreased the expression of VE-PTP at both mRNA and protein levels, suggesting that NCL not only interacts with VE-PTP but also regulates its expression. This highlights the pro-angiogenesis function of NCL and its direct regulatory role on VE-PTP expression, and proves that the *nxhl-*NCL*-*VE-PTP (*ptprb*) signaling pathway is logical and reasonable for angiogenesis.

**Figure 7.**
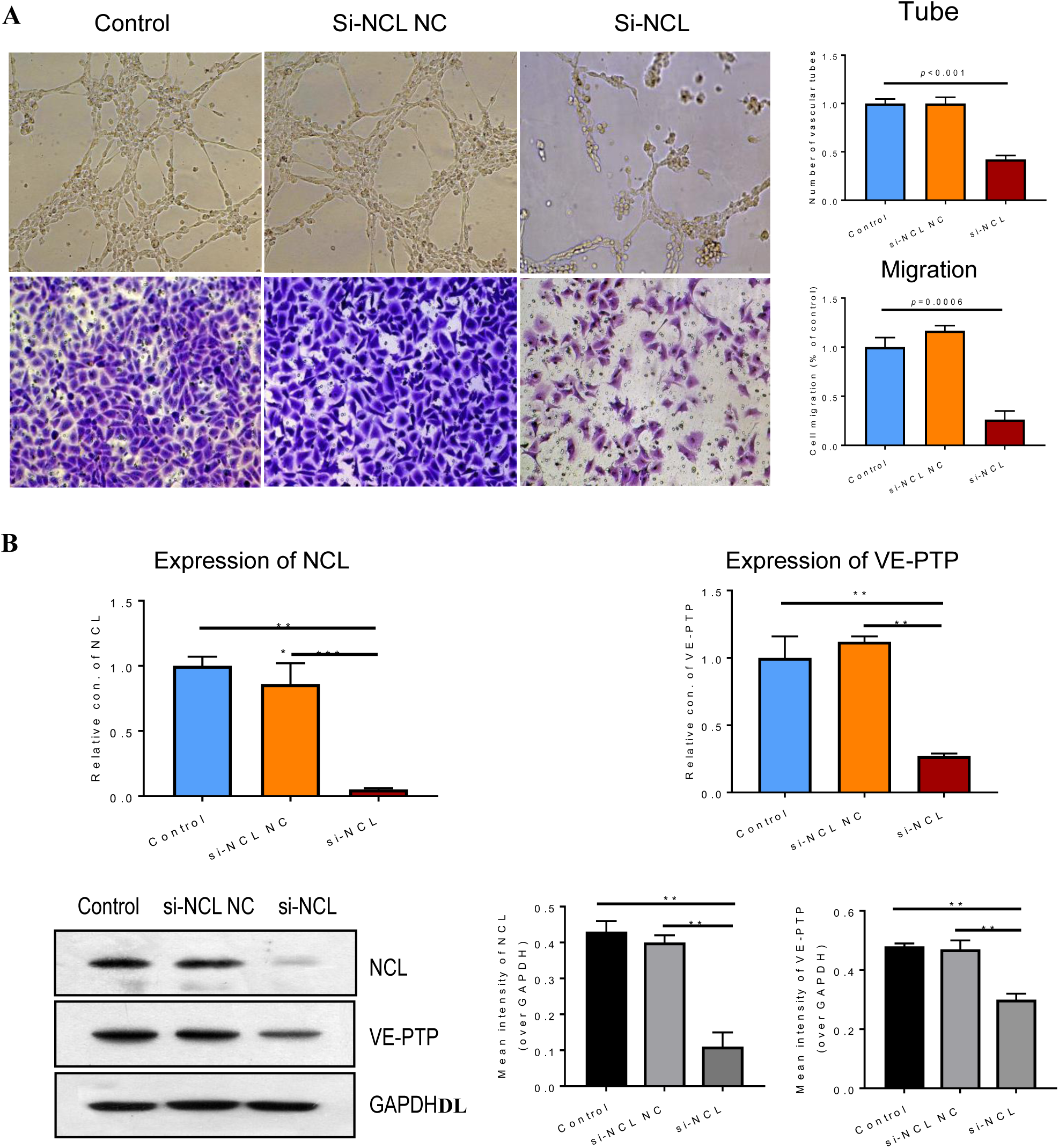
Silence of NCL inhibits angiogenesis and expression of VE-PTP *in vitro*. **A**, Silence of NCL inhibits angiogenesis of HUVECs *in vitro*. The tube formation and cell migration potential of HUVECs treated with si-NCL was determined by using transwell chambers as described in the “Materials and methods” section. Scale bars, 20 μm. Representative images of cells stained in si-NCL treated HUVEC cells. The data represent as mean±SEM from three independent experiments. \**p*<0.05 *p*<0.05, \*\**p*< 0.001 represents statistically significant. **B**, Silence of NCL inhibits the expression of VE-PTP at both mRNA and protein levels. The expression of NCL and VE-PTP was quantified by qPCR. Protein levels of NCL and VE-PTP were examined by using Western blotting post silence of NCL. GAPDH antibody was used as internal control. The gray intensities of the WB images were calculated and present as mean±S EM from three independent experiments. \**p*<0.05 *p*<0.05, \*\**p*< 0.001 represents statistically significant.

### *Nxhl* Controls Angiogenesis by Targeting VE-PTP (*ptprb*)and Linking Angiogenesis Regulatory Genes

It is confirmed that loss of *nxhl* not only downregulates *ptprb* but the angiogenesis landmark genes (Figure 4F), with the addition of finding that *nxhl* binds to NCL which interacts with VE-PTP (*ptprb*), we conclude that *nxhl* controls angiogenesis through *nxhl-*NCL*-*VE-PTP (*ptprb*)-linked angiogenesis regulatory genes. This, for the first time, uncovers the existence of upstream regulatory genes of VE-PTP (*ptprb*). Based on these data, we built a new schematic diagram based on the network in Figure 4F that shows the novel *nxhl-* NCL*-*VE-PTP (*ptprb*) signaling links to the keystone angiogenesis genes (Figure 8A vs. Figure 4F). We also made a schematic diagram to describe the possible mechanism underlying *nxhl-*induced phenotypes of pericardial oedema and vascular patterning defects (Figure 8B). Knockdown of *nxhl* significantly and broadly downregulates angiogenesis-associated landmark genes, including d*ot1L, hand2, erbb2, mef2aa, n2rf1a, hey2, s1pr1, tie2, ptprb, meff2cb, ephB2a, klf2a and cx40*.*8*, through *nxhl-*NCL-VE-PTP (*ptprb*)pathway, while *vegfr2* and *vegfaa* negative feedback control this downregulation. Moreover, loss of *nxhl* increases the phosphorylation of NCL(T76) and decreases the acetylation NCL (K88), indicating that *nxhl* may control angiogenesis by impacting NCL posttranslational modification to regulate downstream VE-PTP signaling pathways. This highlights the crucial role of *nxhl* on angiogenesis development via a hitherto unreported *nxhl-*NCL-VE-PTP (*ptprb*) pathway, which extends the upstream regulatory member of keystone gene VE-PTP (*ptprb*). We conclude that *nxhl* controls angiogenesis by targeting VE-PTP (*ptprb*) through interaction with NCL and linking vascular keystone regulatory genes. Given the extreme importance of the angiogenesis development, and the broad connections with landmark genes, we believe the finding of this novel signaling pathway to be of considerable importance for the study of the angiogenesis development and angiogenesis-dependent diseases.

**Figure 8.**
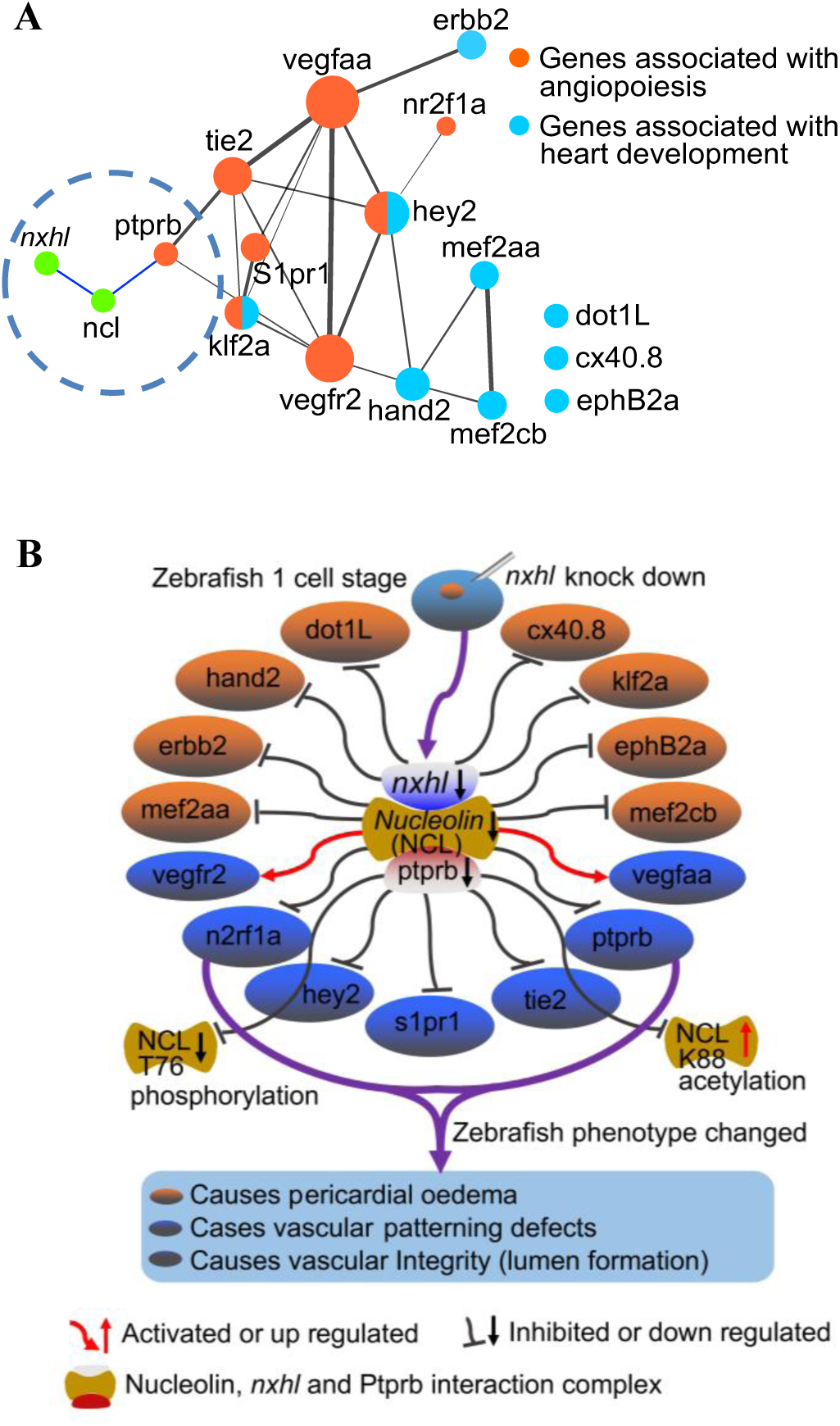
*Nxhl* controls angiogenesis by targeting VE-PTP (*ptprb*)-related angiogenic genes. **A**, Schematic model illustrating the mechanism of *nxhl* in zebrafish angiogenesis and heart development. The interactions between *nxhl* mRNA and NCL protein, NCL protein and *ptprb* mRNA are new-found interactions in this study. **B**, Possible mechanism of *nxhl* in zebrafish angiogenesis and heart development. Knockdown of *nxhl* may downregulate the *nxhl*-NCL-*ptprb* complex, subsequently regulate the proteins associated with angiogenesis and heart development, and finally result in heart pericardial oedema, vascular patterning and integrity defects.

## DISCUSSION

Previous studies showed that VE-PTP is a key player in regulation of angiogenesis and EC adherens junction,^12-15^ and is a potential therapeutic target for angiogenesis-dependent diseases.^7, 8^ It binds to some proteins, such as Tie2, VEGFR2, VE-cadherin and FGD5, that mediate angiogenic signaling pathways.^18, 25-27, 29^ In the present study, we identified a novel zebrafish gene *nxhl*. It controls angiogenic processes *in vitro* and *in vivo*. Deletion of *nxhl* causes angiogenesis-associated phenotypes. Loss of VE-PTP duplicates the phenotypes caused by the upstream *nxhl* deficiency, confirming both act in the same angiogenic signaling pathway. We for the first time show that *nxhl* physically binds to NCL which interacts with VE-PTP and thereby controls angiogenesis. Our study defines a novel *nxhl-*NCL*-*VE-PTP signaling pathway for angiogenesis regulation.

Anti-angiogenic drugs have been a focus of study and lots of inhibitors of angiogenesis are currently used as monotherapy or in combination with chemotherapy or cytokine treatment.^78^ Previous studies showed that AKB-9778, a specific inhibitor of VE-PTP, has demonstrated promising clinic perspective for treatment of angiogenesis-dependent diseases, although it is still under clinical investigation.^31-34^ This highlights the great value of VE-PTP on anti-angiogenic agents. Logically, targeting the upstream regulator of VE-PTP may achieve the same or better effects to that of AKB-9778, because its broader and stronger modulatory forces. However, few upstream regulation mechanisms of VE-PTP has been documented yet. In this study, we identify *nxhl* as a novel powerful upstream regulator of VE-PTP. We find that *nxhl* plays a role in angiogenesis not only because it sharply decreases expressions of VE-PTP and other key angiogenic genes, but also the *nxhl* deletion-caused angiogenesis phenotypes, such as pericardial oedema, defects of caudal fin, intersegmental vessel and caudal vein plexus, are duplicated by the VE-PTP deficiency. These phenotypes of VE-PTP knockdown are mostly identical to a previous study using different morphants to ours.^57^ Additionally, no changes occur in *nxhl* expression upon VE-PTP knockdown, but the expression of VE-PTP significantly decreases upon loss of *nxhl*. This highly implicates that *nxhl* regulates VE-PTP at its upstream and both act in a same signaling pathway. Importantly, the splice-blocking *nxhl* MO displayed phenotypes which are totally overlapping with the translation-blocking MOs, confirming the specificity of phenotypes obtained by *nxhl* injection rather than MO off-target effects. Moreover, *nxhl* controls angiogenesis via ECs migration and tube formation, which is consists with the angiogenic characteristics of VE-PTP on EC adhesion and integrity,^16-18^ confirming its angiogenesis controlling function acts via ECs. This is also strongly supported by our findings that silence of the highly conserve human homologue of *nxhl* not only inhibits the HUVECs migration and tube formation but suppresses the migrations and invasions of cancer cell lines by inhibiting ECs (Figure S9). All these data suggest that *nxhl* is a powerful upstream angiogenesis governor targeting VE-PTP.

On the other hand, the effects of *nxhl* controlling angiogenesis depend on its binding with NCL, which simultaneously bridges *nxhl* and VE-PTP. To our best knowledge, this is the first description on the interactions between *nxhl* and NCL, NCL and VE-PTP, uncovering a novel angiogenesis signaling complex at the upstream of VE-PTP. NCL expresses broadly in all cells in a proliferation-dependent manner ^79^ and almost all compartments of cells. Like VEP-TP, NCL also associates both cancer and other angiogenic diseases. However, this function is more likely related to the cell surface NCL rather than that in other compartments. The cell surface NCL is clustered and highly expressed in ECs of angiogenic blood vessels during angiogenesis,^26, 27, 80^ suggesting that NCL functions as an angiogenic gene. Also, it expresses at the surface of tumor cells, including tumor cells and tumor vasculature. This allows the targeting of different cellular compartments of solid tumors. Additionally, cell surface NCL has been identified in cancer stem cells (CSCs) from different breast cancer cells lines.^81^ Since CSCs are highly tumorigenic,^82, 83^ the association of NCL with the stemness highlights the value of NCL as a potential therapeutic target.^84^ Importantly, dysregulation of NCL associates with higher risk of recurrence or poorer overall survival for some cancers.^85^ These define NCL as both prognostic marker and therapeutic target, highlighting its great value on development of anti-angiogenic drugs.^31, 74^ In this study, we identified the direct interactions between *nxhl* and NCL, and NCL and VE-PTP (*ptprb*) in both zebrafish and 293T cells by ChIRP, RNA Immunoprecipitation and RNA pulldown methods, although we did not yet figure out which subset of NCL (surface, nucleolar or cytoplasmic NCL) participates in this interaction. Importantly, we proved that silence of NCL inhibits angiogenesis of HUVECs and expression of VE-PTP at both mRNA and protein levels. This further supports that NCL plays key roles on angiogenesis by directly controlling downstream VE-PTP. Moreover, deletion of *nxhl* causes a significant decrease of NCL at both mRNA and total protein levels, suggesting that *nxhl* significantly affects and regulates the NCL. This is further supported by the decrease of the phosphorylated T76 and increase of the acetylated K88 of NCL protein upon the *nxhl* knockdown (Figure 6D). Previous study suggested that NCL phosphorylation status heavily affects its cellular compartmentalization.^86^ It promotes EGFR phosphorylation, dimerization and cell growth.^28, 87^ It also promotes HER2 (namely Erbb2) phosphorylation and subsequent MAPK/ Erk pathway activation.^30^ Clinically, combination treatment with NCL and HER2 inhibitors exhibited superior efficacy compared with single treatment in the invasion capacity of breast cancer cells.^88^ Except EGF and HER2, NCL also binds to VEGF,^26^ whose receptor VEGFR2 is tightly associated with VE-PTP resulting in increase of VEGFR2 phosphorylation and activation.^19^ Since EGFR, HER2 and VE-PTP/VEGFR2 have been tightly associated with angiogenesis, we consider *nxhl* may control angiogenesis by affecting NCL phosphorylation which regulates downstream EGFR, HER2 or VE-PTP/VEGFR2 signaling pathways. Notably, in our study, expressions of VEGFR2 and Erbb2 are significantly affected by *nxhl* knockdown, partially supporting this point of view. However, this needs to be investigated in our future works. In addition, NCL acetylation at K88 was previously described *in vivo* and *in vitro*, and this post-translational modification sharply changes its cellular localization. Previous study suggested that NCL may be involved in pre-mRNA synthesis or metabolism because of the presence of NCL-K88ac in nuclear speckles.^89^ However, the characterization and functional significance of NCL acetylation in angiogenesis are still unclear. What the consequences of NCL-K88ac increase on *nxhl* or VE-PTP and subsequent pathways needs further investigation. Taken together, our data enabled us to conclude that *nxhl* regulates the angiogenesis via the *nxhl*-NCL-VE-PTP (*ptprb*) pathway.

The strong power of *nxhl* on angiogenesis controlling also relies on the effects of some other crucial downstream angiogenic genes (such as Tie2, VEGFaa, VEGFR2, S1pr1 and Hey2) which broadly associate VE-PTP signaling (Figure 4, Figure 8). What needs to be stressed is that the expressions of these genes explain the phenotypes induced by the *nxhl* deficiency. They all play irreplaceable roles in multiple aspects of angiogenesis development. For instance, Hand2 is vital in heart development in zebrafish and mouse.^90, 91^ It has been identified as a specifier of outflow tract cells in the mouse by single cell sequencing.^92^ Hey2 mediates the dynamics of cardiac progenitor cells addition to the zebrafish heart.^69^ It is identified as a component of the NKX2-5 cardiac transcriptional network regulating the early stage of the human heart development.^93^ The Dot1L,^63^ Mef2aa,^64^ Mef2cb,^65^ Erbb2,^66^ Klf2a^70^ and EphB2a^94^ also play key roles in the growth of the chamber, cardiomyocyte differentiation, myocardial cell addition, cardiac trabeculation, atrial fibrillation, gap junction, valvulogenesis and myocardial trabeculation during heart development. Importantly, the heart and vascular development are always linked. Previous studies showed that silence of zebrafish S1pr1 not only leads to global and pericardial edema, lack of blood circulation, altered posterior cardinal vein structure, reduced vascularization in ISVs and CVPs,^61^ but also regulates the endothelial barrier integrity via the S1pr1/VE-cadherin/EphB4a pathway.^95^ Similar phenotypes can be induced by the knockdown of Nr2fla in zebrafish due to the decrease of cell proliferation and migration instead of cell death in ECs.^60^ Moreover, mutation of Nr2f1a results in smaller atria due to a specific reduction in the atrial cardiomyocyte number and an increase of the rate of atrial cardiomyocyte differentiation.^96^ Another key gene, Tie2, is essentially required for ISV growth, sprouting, migration, and proliferation of tip cells and acts coordinately with VEGF signaling to control angiogenesis *in vivo*.^97^ Loss of Tie2 leads to death at E10.5 due to vessel remodeling defects and lack of trabeculation.^98^ Notably, Ang-Tie2 system is indispensable for vascular and lymphatic development.^99^ The anti-angiogenic effects of VE-PTP inhibitor, AKB-9778, likely rely on the Ang-Tie2 pathway.^17^ In our study, *nxhl* deletion leads to significant decrease of Tie2, suggesting it regulates not only VE-PTP but also Ang-Tie2 system, which crosstalk with VE-PTP. From this point of review, *nxhl* is a multifunctional master of angiogenesis process. This explains our findings that the phenotypes induced by *nxhl* knockdown mostly resemble the phenotypes caused by deletion of these genes associate with VE-PTP. Although the specific mechanisms underlying need further elucidation, given the extreme importance of these genes in angiogenesis development, we consider the phenotypes caused by *nxhl* morphants as direct or indirect consequences of the down-regulation of VE-PTP and these key genes. We believe the fire-new *nxhl*-NCL-VE-PTP signaling pathway is a highlight for vertebrate angiogenesis development regulation.

In conclusion, we clearly demonstrate that a novel gene *nxhl* controls angiogenesis by targeting VE-PTP through interaction with NCL whose posttranslational modification (phosphorylation and acetylation) may affect downstream VE-PTP signaling pathways. Furthermore, we have elucidated some of the crucial downstream pathways that may be implicated in regulating the angiogenesis. This study reveals a fire-new *nxhl*-NCL-VE-PTP signaling pathway governing vertebrate angiogenesis development, implicating its great potential as therapeutic target for angiogenesis-dependent diseases.

## Acknowledgements

We thank Mr Zhuanbin Wu for zebrafish analysis advices and technical consults.

## Author contributions

H.L.L and X.H.C designed the scientific objectives and oversaw the project. H.L.L., X.H.C., Y.Z.Z., J.X.P., and Y.L discussed the primary ideas of the article. Y.D.Z, J.X.P., Y.L., Y.H., Q.Y.L., P.P.H., C.L.Y., P.Y.W., X.L.C., and P.F.F. collected samples for sequencing DNA and RNA. C.M.J. and their colleagues performed genome sequencing, assembly and annotation. C.M.J. and H.Y.Y performed phylogenomic and whole genome duplication evolution analysis. C.M.J. H.Y.Y., H.L.L., and Y.D.Z performed RNA-seq analysis. H.L.L and Y.D.Z performed functional assay of zebrafish *nxhl* gene and Harbi1 gene. C.M.J, H.Y.Y., H.L.L., and Y.D.Z prepared the supplemental data and method. C.M.J. and H.L.L prepared the draft manuscript with input from all other authors. H.L.L., X.H.C., Y.L., and H.K.Z. discussed and revised the manuscript.

## Data availability

The authors declare that all data reported in this study are fully and freely available from the date of publication. This Whole Genome Shotgun project has been deposited at DDBJ/ENA/GenBank under the accession WOFJ00000000. The version described in this paper is version WOFJ01000000. The draft genome data (genome assembling and annotations) and RNA-Seq data of the embryo are available under BioProject PRJNA574895. Transcriptome (Illumina) data of *nxhl* silence are available in the Sequence Read Archive (SRA) with accession number SRR10199007 and SRR10199008 under BioProject PRJNA573544.

## Supplemental material

Supplemental Information (Notes and Methods)

Supplemental Figures (Figure S1-S16)

Supplemental Tables (Table S1-S28)

Supplemental videos (Movie 1-8)

